# A Polygenic Score for Body Mass Index is Associated with Depressive Symptoms via Early Life Stress: Evidence for gene-environment correlation

**DOI:** 10.1101/536938

**Authors:** Reut Avinun, Ahmad R. Hariri

**Author notes:** Corresponding Author: Reut Avinun, Ph.D., Laboratory of NeuroGenetics, Department of Psychology & Neuroscience, Duke University, Grey Building 2020 West Main St, Ste 0030 Durham, NC 27705.

## Abstract

**Background:** Increasing childhood overweight and obesity rates are associated with not only adverse physical, but also mental health outcomes, including depression. These negative outcomes may be caused and/or exacerbated by the bullying and shaming overweight individuals experience. As body mass index (BMI) can be highly heritable, we hypothesized that a genetic risk toward higher BMI, will predict higher early life stress (ELS), which in turn will predict higher depressive symptoms in adulthood. Such a process will reflect an evocative gene-environment correlation (rGE) wherein an individual’s genetically influenced phenotype evokes a reaction from the environment that subsequently shapes the individual’s health.

**Methods:** We modeled genetic risk using a polygenic score of BMI derived from a recent large GWAS meta-analysis. Self-reports were used for the assessment of ELS and depressive symptoms in adulthood. The discovery sample consisted of 524 non-Hispanic Caucasian university students from the Duke Neurogenetics Study (DNS; 278 women, mean age 19.78±1.23 years) and the independent replication sample consisted of 5 930 white British individuals from the UK biobank (UKB; 3 128 women, mean age 62.66±7.38 years).

**Results:** A significant mediation effect was found in the DNS (indirect effect=.207, bootstrapped SE=.10, 95% CI: .014 to .421), and then replicated in the UKB (indirect effect=.04, bootstrapped SE=.01, 95% CI: .018 to .066). Higher BMI polygenic scores were associated with higher depressive symptoms through the experience of higher ELS.

**Conclusions:** Our findings suggest that evocative rGE may contribute to weight-related mental health problems and stress the need for interventions that aim to reduce weight bias, specifically during childhood.

---

Overweight individuals suffer from stigmatization, bias, and bullying, from multiple sources including peers, health care providers, educators, and, most surprisingly perhaps, family members [1]. In a study of adolescents enrolled in weight loss camps, 37% reported being teased or bullied by a parent [2]. Another study on 2 449 women recruited from a weight loss support group organization, found that 44% experienced stigma from their mothers more than once, while 34% experienced it from their fathers [3]. As weight-related teasing has been shown to predict depression and lower self-esteem [4, 5], it may represent another form of early life stress (ELS) that is associated with various negative physical and mental health outcomes [6, 7].

Gene environment correlations [rGE; 8, 9] can represent passive, evocative, and active processes that create associations between individuals’ genes and the environment. Evocative rGE, which refers to instances in which a genetically influenced phenotype of an individual evokes a certain reaction from the environment, may be relevant to weight-related teasing and bullying, so that individuals with a genetic propensity toward a higher body mass index (BMI), will be more likely to experience teasing, especially in the current Western cultural climate, which is characterized by negative and prejudicial attitudes towards overweight and obese individuals [10].

A recent meta-analysis of genome-wide association studies (GWAS; [11]), consisting of 681 275 participants on average, explained 5% of the variance in BMI with GWAS significant single nucleotide polymorphisms (SNPs). In the current study, we hypothesized that a polygenic score based on the results from this meta-analysis, will predict early life stress (ELS), consistent with evocative rGE, which in turn will predict depressive symptoms in adulthood. We tested our hypothesis in two independent samples: a discovery sample of 524 non-Hispanic Caucasian university students from the Duke Neurogenetics Study and a replication sample of 5 930 adult white British volunteers from the UK Biobank (UKB). Although the GWAS meta-analysis included data from the UKB, current BMI was not a phenotype of interest in our study, and therefore the overlap should not bias our analyses. Nonetheless, to validate our results in the analyses that included UKB data, we also used BMI polygenic scores that were based on a GWAS that did not include the UKB as a discovery sample [12].

## MATERIALS AND METHODS

### Participants

Our discovery sample consisted of 524 self-reported non-Hispanic Caucasian participants (278 women, mean age 19.78±1.23 years) from the Duke Neurogenetics Study (DNS) for whom there was complete data on genotypes, ELS, depressive symptoms, and all covariates. All procedures were approved by the Institutional Review Board of the Duke University Medical Center, and participants provided informed consent before study initiation. All participants were free of the following study exclusions: 1) medical diagnoses of cancer, stroke, diabetes requiring insulin treatment, chronic kidney or liver disease, or lifetime history of psychotic symptoms; 2) use of psychotropic, glucocorticoid, or hypolipidemic medication; and 3) conditions affecting cerebral blood flow and metabolism (e.g., hypertension). Importantly, neither current nor lifetime diagnosis were an exclusion criterion, as the DNS sought to establish broad variability in multiple behavioral phenotypes related to psychopathology.

The replication sample consisted of 5 930 white British individuals (3 128 women, mean age 62.66±7.38 years), who participated in the UKB’s imaging wave, completed an online mental health questionnaire [13], and had complete genotype, ELS, depressive symptoms and covariate data. The UKB [www.ukbiobank.ac.uk; 14] includes over 500,000 participants, between the ages of 40 and 69 years, who were recruited within the UK between 2006 and 2010. The UKB study has been approved by the National Health Service Research Ethics Service (reference: 11/NW/0382), and our analyses were conducted under UKB application 28174.

### Race/Ethnicity

Because self-reported race and ethnicity are not always an accurate reflection of genetic ancestry, an analysis of identity by state of whole-genome SNPs in the DNS was performed in PLINK [15]. The first two multidimensional scaling components within the non-Hispanic Caucasian subgroup were used as covariates in analyses of data from the DNS. The decision to use only the first two components was based on an examination of a scree plot of the variance explained by each component. For analyses of data from the UKB, only those who were ‘white British’ based on both self-identification and a principal components analysis of genetic ancestry were included. Additionally, the first 10 multidimensional scaling components received from the UKB’s data repository (unique data identifiers: 22009-0.1-22009-0.10) were included as covariates as previously done [e.g., 16].

### Body Mass Index (BMI)

In both DNS and UKB samples, BMI was calculated at the time of imaging based on the height and weight of the participants. In the DNS, this calculation was based on imperial system values (pounds/inches2*703), while in the UKB the metric system was used (kg/m2). In the DNS 1.3% of the sample was obese, compared to 18.7% in the UKB.

### Depressive symptoms

In the DNS, the 20-item Center for Epidemiologic Studies Depression Scale (CES-D) was used to asses depressive symptoms in the past week [17]. All items were summed to create a total depressive symptoms score. In the UKB, the Patient Health Questionnaire 9-question version (PHQ-9) was used to asses depressive symptoms in the past 2 weeks [18]. All items were summed to create a total depressive symptoms score.

### Early life stress

In the DNS, ELS was estimated using the Childhood Trauma Questionnaire [CTQ; 19]. The CTQ has 28-items and it assesses the frequency of emotional, physical, and sexual abuse as well as emotional and physical neglect. The scores on the 5 subscales (each ranging from 5 to 25) were summed to create a total score of ELS. In the UKB, the Childhood Trauma Screener – 5 item (CTS-5) was used to assess adverse events during childhood [20]. CTS-5 is a short version of the CTQ consisting of 5 items: “Felt hated by family member as a child”, “Physically abused by family as a child”, “Felt loved as a child” (reverse coded), “Sexually molested as a child”, and “Someone to take to doctor when needed as a child” (reverse coded). The 5 items, each ranging from 0-4, were summed to create a total score of ELS.

### Genotyping

In the DNS, DNA was isolated from saliva using Oragene DNA self-collection kits (DNA Genotek) customized for 23andMe (www.23andme.com). DNA extraction and genotyping were performed through 23andMe by the National Genetics Institute (NGI), a CLIA-certified clinical laboratory and subsidiary of Laboratory Corporation of America. One of two different Illumina arrays with custom content was used to provide genome-wide SNP data, the HumanOmniExpress (N=329) or HumanOmniExpress-24 [N=195; 21, 22, 23]. In the UKB, samples were genotyped using either the UK BiLEVE (N=569) or the UKB axiom (N=5,361) array. Details regarding the UKB’s quality control can be found elsewhere[24].

### Quality control and polygenic scoring

For genetic data from both the DNS and UK Bionbank, PLINK v1.90 [15] was used to apply quality control cutoffs and exclude SNPs or individuals based on the following criteria: missing genotype rate per individual >.10, missing rate per SNP >.10, minor allele frequency <.01, and Hardy-Weinberg equilibrium p<1e-6. Additionally, in the UKB, quality control variables that were provided with the dataset were used to exclude participants based on a sex mismatch (genetic sex different from reported sex), a genetic relationship to another participant, outliers for heterozygosity or missingness (unique Data Identifier 22010-0.0), and UKBiLEVE genotype quality control for samples (unique Data Identifiers 22050-0.0-22052-0.0).

Polygenic scores were calculated using PLINK’s [15] “--score” command based on published SNP-level summary statistics from a recent BMI GWAS meta-analysis [11]. SNPs from the GWAS of BMI meta-analysis were matched with SNPs from the DNS and the UKB. For each SNP the number of the alleles (0, 1, or 2) associated with BMI was multiplied by the effect estimated in the GWAS. The polygenic score for each individual was an average of weighted BMI-associated alleles. All SNPs matched with SNPs from the DNS and UKB were used regardless of effect size and significance in the original GWAS, as previously recommended and shown to be effective [25, 26]. A total of 442 040 SNPs from the DNS and 648 530 SNPs from the UKB were included in the polygenic scores. The approach described here for the calculation of the polygenic scores was successfully used in previous studies [e.g., 27, 28-30]. For validation of the indirect effect in the UKB, BMI polygenic scores were also calculated based on an older GWAS that did not include the UKB as a discovery sample [12].

### Statistical analysis

Linear regression analyses in SPSS v25 were conducted to test for an association between the BMI polygenic score and BMI in adulthood. The PROCESS SPSS macro, version 3.1 [31], was used to conduct the mediation analyses. Participants’ sex (coded as 0=males, 1=females), age, and ethnicity genomic components were entered as covariates in all analyses. In the mediation analyses, bias-corrected bootstrapping (set to 5,000) was used to allow for non-symmetric 95% confidence intervals (CIs). Specifically, indirect effects are likely to have a non-normal distribution, and consequently the use of non-symmetric CIs for the determination of significance is recommended [32]. However, bias-corrected bootstrapping also has its faults [33] and, consequently, as supportive evidence for the indirect effect, we also present the test of joint significance, which examines whether the *a path* (BMI polygenic score to ELS) and the *b path* (ELS to depressive symptoms, while controlling for the BMI polygenic score) are significant. The BMI polygenic scores were standardized to make interpretability easier. The mediation was first analyzed in the DNS, and then a replication was tested in the UKB. As a validation of the indirect effect in the UKB, it was also tested with an older BMI polygenic score that was not based on a GWAS that included the UKB [12].

## RESULTS

Descriptive statistics are presented in table 1.

**Table 1.** Descriptive statistics of study variables.

|  | DNS |  |  |  | UK Biobank |  |  |  |
| --- | --- | --- | --- | --- | --- | --- | --- | --- |
|  | Min | Max | Mean | SD | Min | Max | Mean | SD |
| **Age** | 18 | 22 | 19.78 | 1.24 | 45 | 78 | 62.66 | 7.38 |
| **BMI** | 16.30 | 39.15 | 22.29 | 2.83 | 14.94 | 58.04 | 26.60 | 4.419 |
| **Early life stress** | 25 | 74 | 31.29 | 7.16 | 0 | 20 | 1.68 | 2.32 |
| **Depressive symptoms** | 0 | 43 | 8.99 | 7.18 | 0 | 27 | 2.45 | 3.39 |

### Confirming an association between BMI polygenic scores and measured BMI

As a preliminary analysis we confirmed that higher BMI polygenic scores were significantly associated with higher measured BMI in both the DNS (N=522, b=.837, SE=.117, p<.001) and the UKB (N=5 925, b=1.41, SE=.054, p<.001). (notably, the UKB was included in the BMI GWAS, and consequently the significant association is expected and possibly somewhat inflated). These associations were robust to the inclusion of sex, age, and ethnicity genomic components as covariates. The sample sizes for these analyses were slightly different from the mediation analyses below because measured BMI was missing for a few participants.

### BMI polygenic scores predict ELS (a path) in the DNS

The BMI polygenic scores were significantly associated with ELS (b=.65, SE=.31, p=.038), so that higher scores predicted higher ELS. Of the covariates, age was significantly and negatively associated with ELS (b=-.73, SE=.25, p<.01).

### ELS predicts depressive symptoms (b path) in the DNS

With the BMI polygenic scores in the model, ELS significantly and positively predicted depressive symptoms (b=.32, SE=.04, p<.001).

### BMI polygenic scores predict depressive symptoms in the DNS

The BMI polygenic scores did not significantly predict depressive symptoms (b=-.34, SE=.31, *ns*). Notably, however, the significance of a direct path from X (BMI polygenic scores) to Y (depressive symptoms) or the ‘total effect’ (the ‘c’ path), is not a prerequisite for the testing of a mediation/indirect effect [34-36].

### Mediation model in the DNS

The indirect path (*a*b*), BMI polygenic scores to ELS to depressive symptoms was significant as indicated by the bias corrected bootstrapped 95% CI not including zero (Figure 1a; indirect effect=.207, bootstrapped SE=.10, 95% CI: .014 to .421).

**Figure 1.**
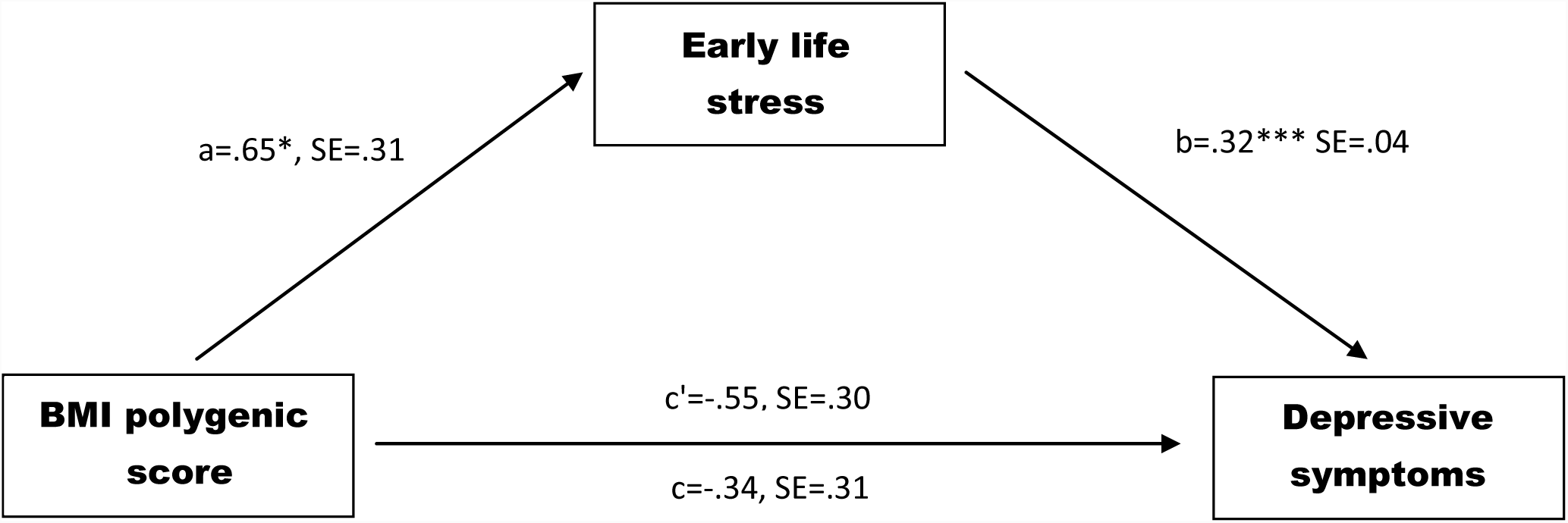
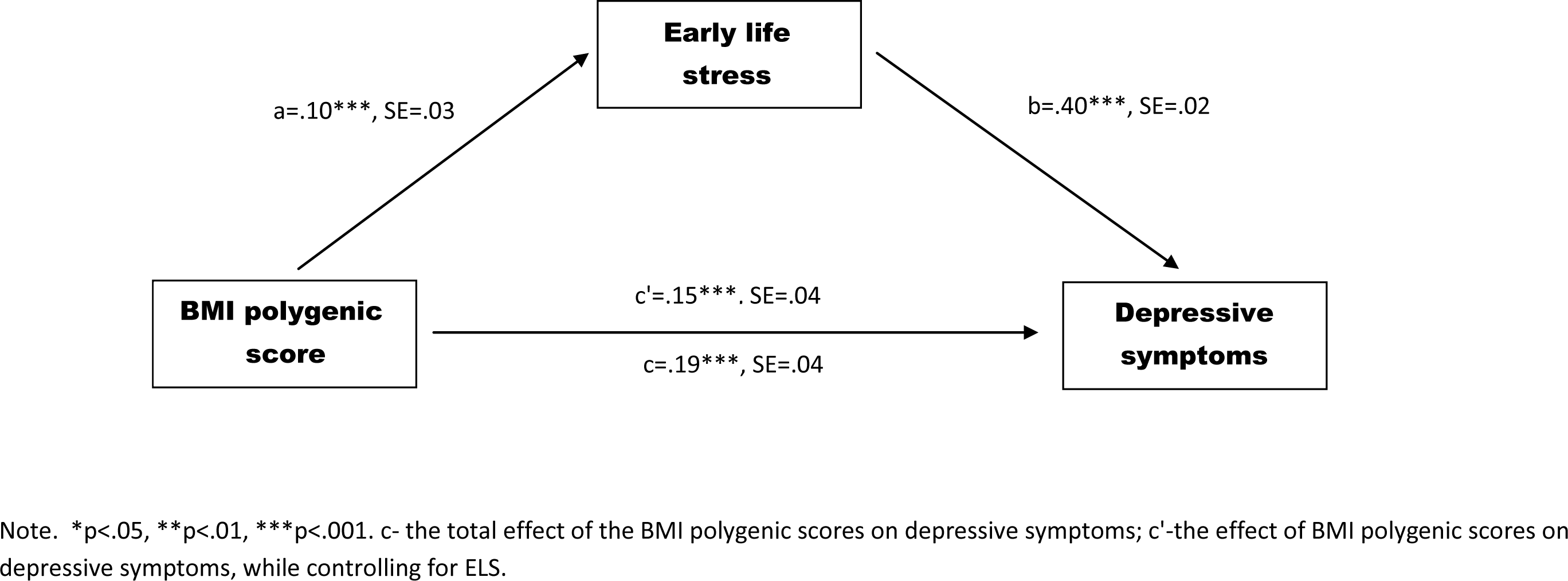
Mediation model linking genetic risk for higher BMI to higher depressive symptoms, via elevated levels of early life stress **1a.** Duke Neurogenetics Study: Discovery sample **1b.** UK Biobank: Replication sample

### Mediation Model in the UBK

The *a path*, from the BMI polygenic scores to ELS, and the *b path*, from ELS to depressive symptoms while controlling for BMI polygenic scores, were significant (*a path*: b=.10, SE=.03, p<.01; *b path*: b=.40, SE=.02, p<.001). The indirect path also replicated (Figure 1b; indirect effect=.04, bootstrapped SE=.01, 95% CI:.018 to.066), supporting a mediation in which BMI polygenic scores are associated with depressive symptoms indirectly through ELS. Similar results were obtained with the BMI polygenic scores that were based on a GWAS that did not include the UKB as a discovery sample (indirect effect=.026, bootstrapped SE=.01, 95% CI: .004 to .05).

## DISCUSSION

Here, in two independent samples, we provide novel evidence supporting evocative rGE as a possible mechanism in weight-related depression. We demonstrate a significant mediation in which higher GWAS-derived BMI polygenic scores are associated with higher levels of depressive symptoms in adulthood through elevated levels of ELS. These results suggest that in the current Western cultural climate, having a genetic makeup that increases the risk of a high BMI, may lead to a phenotype that evokes increased stress, which increases the experience of depressive symptoms in adulthood.

Various studies have reported links between being overweight and experiencing stigmatization, teasing, and bullying from peers, educators, co-workers, health care providers, and family members [1]. This negativity can lead to adverse mental health outcomes, including depression [5], but is not limited to mental health. Obesity, childhood trauma, and depression have all been linked to physical illness including cardiovascular disease, type 2 diabetes, and autoimmune disorders [6, 37-40].

While several strategies have been proposed to battle the growing prevalence of childhood obesity, including nutrition standards for school meals; improved early care and education; and increased access to adolescent bariatric surgery [41], our findings further encourage weight stigma reduction efforts, specifically among family members and parents. In addition to the myriad of mental and physical health disorders that are associated with ELS and childhood trauma, one of the most prevalent coping responses to weight stigma is eating [3]. Consequently, ELS may lead to additional weight gain and is itself a risk factor for obesity [42]. Thus, interventions that aim to reduce weight stigma may have a broad positive effect on health.

Although our study has several strengths, including the use of two independent samples with markedly different characteristics (e.g., young university students versus older community volunteers) and a GWAS-derived polygenic score, it is not without limitations. First, retrospective reports were used for the estimation of ELS and childhood trauma. Ideally, prospective data should be used to model ELS in the absence of reporting bias. Second, we did not have measures of childhood BMI in either sample. Although previous research does support a link between childhood BMI, teasing, and depression, and genetic influences on BMI have been shown to be relatively stable throughout development [43, 44], genetically informed longitudinal studies across development are needed to further validate our findings. Third, the non-Hispanic Caucasian DNS sample is relatively homogeneous in terms of social background, which may have led to an underestimation of the effect in this sample. Fourth, our findings are limited to populations of European descent and to the Western culture. Additional research in diverse populations is needed to determine the extent to which the observed evocative rGE mechanism shapes weight-related mental health. Further replication is also needed to evaluate the potential of the BMI polygenic score as a risk biomarker of depression associated with ELS.

